# Observation of tandem running behavior in mating pairs of Asian dampwood termite, *Hodotermopsis sjostedti*

**DOI:** 10.1101/2025.05.02.651862

**Authors:** Nobuaki Mizumoto, William Chambliss, Carroll P Elijah, Nakazono Tomohiro, Taisuke Kanao

## Abstract

As a social insect, termite colonies can grow to a group of millions of individuals, yet all colonies start from a single mating pair. Recent studies indicate that the pair formation process shows a large diversity among species, especially in basal lineages. Thus, comparative information is integral to estimating the ancestral state of this essential stage of the termite life cycle. The Asian dampwood termite, *Hodotermopsis sjostedti*, has been well-studied as a model basal termite of caste differentiation processes. Yet, their pair formation remains undocumented. In this study, we found that mating pairs of *H. sjostedti* show clear tandem running behavior. Both females and males played a leading role, with females showing more leader roles, and they switched their leading roles even within the same pair. We also found that dish size affected tandem movement coordination; pairs showed faster and more stable tandem running in a larger dish. We provide a tracking dataset of 17 body parts, including antennal and leg movements during tandem runs, which can be utilized in future comparative studies. This study supports the idea that tandem running existed in the early ancestors of termites and sheds light on the origin of termite mate pairing.

## Introduction

Social insects play a dominant role in ecosystems, either as predators, pollinators, or decomposers, contributing to global biomass (Bar-On et al., 2018; Eggleton, 2020; Tuma et al., 2020). The ecological success of social insects is often owed to the large size of their colony, ranging from hundreds to millions of individuals. Thus, extensive research efforts have focused on colony functions, regulated by their caste systems, where parents monopolize reproduction, and offspring will either develop into working castes that are responsible for colony tasks or alates that disperse to start a new colony (Noirot, 1991; Oster and Wilson, 1978). However, highlighting mature colonies of social insects often obscures the fact that most colonies need to start from one or a few reproductive individuals dispersed from their original colonies, except for a few species (Cronin et al., 2013). The first critical task of these dispersers is finding a mating partner; such pairing behavior is as important as sophisticated social behaviors to complete their colony life cycles.

Termites are one of the major lineages of eusocial insects and have evolved from subsocial wood-feeding cockroach ancestors (Bell et al., 2007). Termite colonies usually start with a monogamous mating pair, which will be a king and a queen in the mature colony (Chouvenc, 2022; Nutting, 1969). Termite mate pairing is often described as follows: in a short period of the year, numerous alates fly off to disperse. Once they land on the ground, they shed their wings to walk to search for a mating partner. Upon encounter, a pair performs tandem running, with the males following the females while searching for a nest site. However, this description is biased toward the observation of several neoisopteran termites, and pairing processes are documented to be more diverse, especially in other lineages (Mizumoto et al., 2022). Some do not show tandem running, but females and males separately come to the nest sites (Sugio et al., 2020; Wilkinson, 1962). Some show tandem running, but the leader role is more flexible (Grasse, 1942; Lüscher, 1951; Mizumoto et al., 2022). Furthermore, *Cryptocercus* woodroach, a sister group of termites, should adopt a distinct pairing process from termites, as they are socially monogamous but genetically not (Yaguchi et al., 2021). Therefore, it is important to study the diversity of tandem running behavior, especially in basal lineages, which are often cryptic.

Asian dampwood termite, *Hodotermopsis sjostedti*, belongs to Hodotermopsidae, one of the early diverging groups of termites (Wang et al., 2022). This species is extensively studied for the caste development system (e.g., (Kobayashi et al., 2023; Koshikawa et al., 2005; Miura et al., 2004, 2000; Nii et al., 2019; Oguchi et al., 2016; Oguchi and Miura, 2023; Shimoji et al., 2019)). However, their basic biology is not well understood. For example, termite nesting strategies can be classified based on how they utilize their food and nest resources (Abe, 1987; Korb, 2008; Mizumoto and Bourguignon, 2020), and *H. sjostedti* was originally classified as a one-piece nester whose entire colony is completed within a single piece of wood (Abe, 1987). However, a later field study clearly demonstrated that this species is actually a multiple-piece nester that nests across multiple wood pieces by interconnecting them with underground tunnels (Kitade et al., 2012). In terms of mate pairing, there are several studies on swarming flight in nature (Ohmura and Makihara, 2005), developmental mechanisms (Kobayashi et al., 2023; Miura et al., 2000; Oguchi et al., 2016; Oguchi and Miura, 2023), and identification of sex-specific chemicals (Lacey et al., 2011), yet no information about the mate pairing process has been documented for this species.

Here, we study the tandem running behavior of *H. sjostedti*. We observe their tandem running in the same methodological framework as previous studies in other genera (Mizumoto et al., 2021; Mizumoto and Dobata, 2019). Also, we qualify their behavior using deep learning posture tracking to compare the female leader and the male leader. Finally, given the large body size of this species, we compare the observation between two different-sized dishes.

## Methods

### Behavioral observation

The colony of *Hodotermopsis sjostedti* was collected at Yakushima Island, Kagoshima prefecture, on 12 May 2023. The colony included nymphs, which differentiated into alates in a few months. In the laboratory, the colony was separated into subgroups and maintained in plastic containers under dark conditions, either with the brown-rotted pinewood mixed cellulose (BPC) (Mitaka et al., 2023) and blocks of pinewood under 25°C, or with original nesting wood, commercial sawdust mat developed for rearing rhinoceros beetles, and blocks of pinewood under 23°C. In July 2023, both subgroups produced alates. We moved the plastic containers with nests to 27°C, and alates flew to disperse from the nests. Alates were then collected, separated by sex, introduced to be dealated by manually pinching wings with forceps, and color-marked with one dot of paint (PX-20; Mitsubishi) on the abdomen to distinguish sex identities (Fig. 1A). These termites were isolated for at least 30 minutes before the experiments.

**Figure 1.**
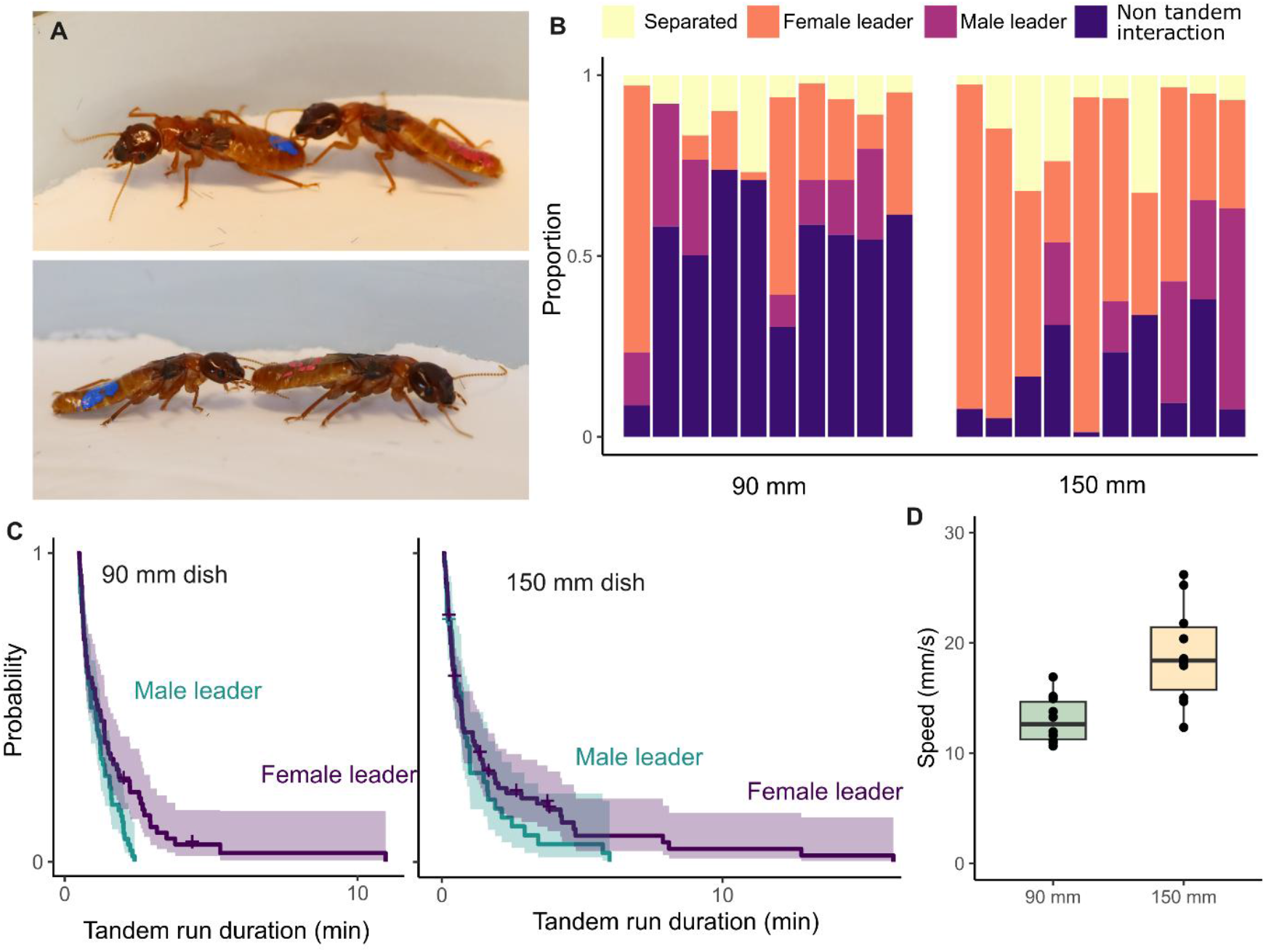
Description of tandem running behavior in *Hodotermopsis sjostedti*. (A) Male-led and female-led tandem runs. Termites with red markings are females and those with blue markings are males. (B) Proportion of time in each state of the pair during observation. Each bar represents one pair. (C) Comparison of tandem running between female-led and male-led tandem running, and two different sizes of experimental arena. Kaplan–Meier survival curves were generated for each species, and the symbol “+” indicates the censoring due to the end of observations. (D) Comparison of leader moving speed during tandem running between different sizes of dish.

We introduced a female-male pair into the experimental arena. The experimental arenas consist of a petri dish (φ = 87 and 143 mm, respectively; hereafter, we call these 90 and 150 mm, as these are the manufactured sizes with lids) covered with a layer of moistened plaster that was polished before each trial. We recorded termite movements in the arena for 30 minutes using a video camera (HC-X1500-K, Panasonic) with a resolution of 3840×2160 pixels at 59.96 frames per second (FPS). Some of the experimental replicates escaped from the arena; these replicates were cropped when an individual first exited the arena (data include replicates with 12, 15, 24, and 28 min in 150 mm arena). In total, we obtained videos of 10 pairs in the 90 mm dish and 11 pairs in the 150 mm dish. Because only one colony was available, all pairs were nestmates. All the videos were cropped to 2000×2000 pix to only include the arena in the frame before the video analysis.

### Data processing

All videos were analyzed using SLEAP v 1.4.0 (Pereira et al., 2022) to estimate the movement of the body parts of each individual. The model was based on that developed for *Reticulitermes speratus* and *Coptotermes formosanus* in a previous study (Mizumoto and Reiter, 2025), with a 17-node skeleton: antenna tips (left and right: LR), antenna middle (LR), antenna base (LR), head (middle of mouth parts), head-pronotum boundary, pronotum-mesonotum boundary, metanotum-abdomen boundary, abdomen-tip, fore legs (LR), mid legs (LR), and the hind legs (LR). First, we tracked the videos at a 90 mm dish. We labeled 213 frames (426 instances) across 20 individuals from 10 videos for training. We trained a U-Net-based model with a multi-animal top-down approach, with a receptive field size of 316 pixels for the centroid and the centered instance, on Nvidia GeForce RTX 4090, where augmentation was done by rotating images from -180 to 180 degrees. The mean Average Precisions (mAP) of the centroid model and instance model were 0.35 and 0.76, respectively. While tracking after the inference, we used the instance similarity method with the greedy matching method. Then, based on the model developed for the 90 mm dish, we performed the tracking for the 150 mm dish. We used the same procedure as above, where we labeled 22 frames (44 instances) across 18 individuals from 9 videos for training.

All pose estimation data were converted to HDF5 files, which were further converted into FEATHER files using Python. We employed a linear interpolation method to address missing values in the dataset and used a median filter with a kernel size of 5 to reduce noise. We extracted the position of the head, body center (metanotum-abdomen boundary), and abdomen-tip for further analysis.

### Data analysis

All data analysis was performed in R v. 4.4.3 (R Core Team, 2024). We downsampled the data into 5 FPS and scaled all data from pixels to mm (2000 pixels = arena size). We automatically determined whether pairs were in tandem and who was the leader for each frame, based on the postures and spatial position of partners. Tandem running was defined as a state in which both individuals were (1) in proximity, (2) moving above a minimum speed threshold, and (3) aligned in heading direction. Specifically, proximity was determined by measuring the Euclidean distance between the body centers of the two termites; pairs were considered “close” when this distance was less than 0.6 times the sum of their body lengths. The heading of each termite was calculated as a vector from the tip of the abdomen to the head tip, and the absolute angular difference in heading was used to evaluate alignment. Tandem running was defined when the absolute heading difference was below 60 degrees (i.e., π/3 radians), and both individuals exceeded a movement threshold of 1.213 mm/s, based on the first quartile of female speed distribution. To avoid misclassifying brief interactions as true tandem runs, tandem run classifications were smoothed using a run-length encoding (RLE) based filter. Tandem or separation segments shorter than 4 seconds (20 frames at 5 FPS) were merged with adjacent longer segments. Leader roles were determined within tandem runs by comparing relative distances between the leader’s head and the follower’s abdomen. Specifically, if the distance from the male’s head to the female’s abdomen exceeded the reverse (female’s head to male’s abdomen), the male was considered to be leading; otherwise, the female was considered the leader. These frame-wise leader states were further smoothed using the same RLE-based filter, requiring at least 3 seconds (15 frames) of consecutive leadership to qualify as a valid leader period. Switching events were defined as changes in the leader identity during tandem runs. A switch was counted only when a change in leader identity occurred between consecutive frames classified as tandem. Switching frequency was computed for each video, and the number of role changes per tandem period was recorded. The thresholds used during these classification processes were determined arbitrarily based on the video inspection. Minor quantitative changes in these did not affect our conclusion.

We compared the duration of each tandem running event between leader sexes and two different dish sizes, using mixed-effects Cox models, with leader sex, dish size, and their interaction being treated as a fixed effect and each pair id as a random effect. We used the coxme() function in the coxme package in R (Therneau, 2015). The likelihood ratio test was used to test for statistical significance of the explanatory variable (type II test), with the Anova() function in the car package. We also compared the number of leader switching events between two different dish sizes, using a generalized linear model with Poisson error and a log link function. Finally, we compare the moving speed of leaders between the two dish sizes using a t-test.

## Results

The termite, *Hodotermopsis sjostedti*, showed a clear tandem running behavior with both females and males performing leader roles (Fig. 1A, Video 1). We also observed both female-female and male-male same sex tandem running during sample preparation, as in the other termite species (Matsuura et al., 2002; Mizumoto et al., 2022, 2024b). We detected tandem running behavior from all the pairs we observed (Fig. 1B), but the patterns were different between leader sexes and dish sizes. In general, we observed more frequent female-led tandem runs than male-led tandem runs (Fig. 1B), where female-led tandems were more stable than male-led tandems (mixed-effects Cox model, χ^2^1 = 8.69, *P* = 0.003; Fig. 1C). The pattern was consistent between two different dish sizes (dish size: χ^2^1 = 2.17, *P* = 0.14, interaction: χ^2^1 = 0.66, P = 0.416; Fig. 1C). Leader role was swapped even within a pair during 30 minute observations (Table 1), with no difference between dish sizes (GLM, χ^2^1 = 0.067, *P* = 0.796).

**Table 1.** The number of leader role switches for each pair.

| video | dish size | leader switches |
| --- | --- | --- |
| Hod_sjo_FM_90_A_02 | 90 mm | 0 |
| Hod_sjo_FM_90_A_03 | 90 mm | 2 |
| Hod_sjo_FM_90_A_05 | 90 mm | 1 |
| Hod_sjo_FM_90_A_06 | 90 mm | 2 |
| Hod_sjo_FM_90_A_07 | 90 mm | 1 |
| Hod_sjo_FM_90_A_08 | 90 mm | 0 |
| Hod_sjo_FM_90_A_09 | 90 mm | 0 |
| Hod_sjo_FM_90_A_10 | 90 mm | 1 |
| Hod_sjo_FM_90_A_14 | 90 mm | 0 |
| Hod_sjo_FM_90_A_15 | 90 mm | 1 |
| Hod_sjo_FM_150_A_02 | 150 mm | 1 |
| Hod_sjo_FM_150_A_03 | 150 mm | 2 |
| Hod_sjo_FM_150_A_05 | 150 mm | 1 |
| Hod_sjo_FM_150_A_06 | 150 mm | 0 |
| Hod_sjo_FM_150_A_07 | 150 mm | 2 |
| Hod_sjo_FM_150_A_08 | 150 mm | 0 |
| Hod_sjo_FM_150_A_09 | 150 mm | 1 |
| Hod_sjo_FM_150_A_10 | 150 mm | 0 |
| Hod_sjo_FM_150_A_14 | 150 mm | 0 |
| Hod_sjo_FM_150_A_15 | 150 mm | 0 |

During tandem running, the leader movement speeds were influenced by the dish size, where termites moved faster in a larger dish (t-test, *t*_12.85_ = 3.82, *P* = 0.002; Fig. 1D).

## Discussion

Our observations clearly showed that dealates of *Hodotermopsis sjostedti* exhibit tandem running behavior, with both females and males playing both leader and follower roles (Fig. 1, Video 1). In many species of termites, females play a leader role in tandem running (Mizumoto et al., 2022). Still, documentation of tandem running with both female and male leaders has been limited in several Kalotermitidae species (summarized in (Mizumoto et al., 2022)), including *Kalotermes flavicollis* (Grasse, 1942; Lüscher, 1951), *Cryptotermes havilandi* (Lüscher, 1951) (but suspected to be *C. dudleyi*. See (Mizumoto et al., 2022)), *Paraneotermes simplicicornis* (Carr, 1972), and *Glyptotermes fuscus* and *G. satsumensis* (Mizumoto et al., 2022). In addition, the fossil record indicates that the extinct kalotermitid termite, *Electrotermes affinis*, shows male-led tandem running behavior (Mizumoto et al., 2024a). However, even though the previous ancestral state reconstruction estimated that the ancestor of termites exhibited tandem running, with both females and males being leaders (Mizumoto et al., 2022), there was no record of that behavior in other lineages than Kalotermitidae. Thus, our observation on *H. sjostedti* is critical as this species belongs to the family Hodotermopsidae which is part of the distinct clade Teletisoptera comprising Hodotermopsidae, Stolotermitidae, Hodotermitidae, and Archotermopsidae. As an early diverging lineage, the clade Teletisoptera separated from other Euisoptera 117.9 million years ago (Wang et al., 2022), providing the informative piece to infer the evolutionary origin of tandem running behavior.

Although the biology of *Hodotermopsis* is often compared with that of *Zootermopsis* as a related species (e.g., (Miura et al., 2004)), our observations suggest that these two groups use distinct mate-pairing processes. The pairing process of *Zootermopsis* species has been documented in several papers (Castle, 1934; Howse, 1970; Shellman-Reeve, 2001, 1999, 1994; Stuart, 1969), their use of tandem running behavior is less clear (summarized in Fig. S12 in (Mizumoto et al., 2022)). A previous study treated the tandem running status of *Zootermopsis* as a female-led tandem (Mizumoto et al., 2022), yet original descriptions clearly mention that the tandem pairing is weaker than other termite species showing tandem running behavior (Castle, 1934; Howse, 1970). By using the same experimental setup as the current study, we could not observe the clear tandem running in *Zootermopsis nevadensis*, collected in Hyogo Prefecture, Japan (although small sample size; *n* = 3 pairs, on Jan 4th, 2022). These observations indicate that the pairing process of *H. sjostedti* is distinct from *Zootermopsis* species. It is reasonable that these species exhibit different nesting types, with *Hodotermopsis* being a multiple-piece nester and *Zootermopsis* being a one-piece nester. One future direction is to study the relationship between nesting habitat and the pairing process in a phylogenetic comparative framework.

One limitation of the current study is that our observation is limited to one colony. Thus, all pairs are inevitably nest-mate pairs, which may have affected the tandem running behavior. Also, in other species, it is known that tandem running behavior can be affected by individual conditions, such as body size (Husseneder and Simms, 2008; Matsuura et al., 2002) and time after swarming (Mizumoto et al., 2024c). Thus, reflecting individual status, there should be a quantitative variation of tandem running propensity across different colonies. However, it is unrealistic to suppose the colony variation in the pairing mode, with some colonies exhibiting tandem running while others using different pairing methods. For example, in *Marginitermes hubbardi*, a laboratory observation demonstrates that this species does not usually show tandem running behavior except for one pair (Carr, 1972). One of the authors observed a tandem running behavior of *M. hubbardi* on the tree trunk in the field condition (one personal observation by N. Mizumoto on July 31, 2019, in Tempe, Arizona), implying that there might be a specific condition for this species to exhibit tandem running behavior. Thus, it might be difficult to prove the lack of tandem running only from the laboratory observations. However, even with limitations, our study provides a positive observation of the clear tandem running behavior of *H. sjostedti*, which should be valid in field environments.

Notably, we found that the termite tandem running behavior may be affected by the size of observational arenas (Fig. 1), with termites moving fast and showing more tandems in the larger arena. Instead, in a smaller arena, pairs of *H. sjostedti* spent more time on non-tandem running interactions, such as grooming. The termite, *H. sjostedti*, is one of the largest termites (12-13 mm body length of dealates in our study) (Mizumoto and Bourguignon, 2021). Since tandem running behavior is an exploratory behavior for a nest site for colony foundation, the 90 mm arena may have been too small to be recognized as an open space for this species. This is in contrast to other smaller species with clear tandem running in 90 mm or even smaller dish sizes (Mizumoto and Reiter, 2025). Because arena size can affect the free walking behavior in insects (Scharf et al., 2024), it could be important to provide a large enough arena for the focal species to exhibit their tandem running behavior.

In conclusion, our study contributes to the understanding of the diversity and evolution of mate-pairing behavior in termites. Even though mate pairing plays a crucial role in the life cycle of termites, little attention has been paid to it compared to other social behaviors. One challenge is that mate pairing is a seasonal event, which can be observed in a limited period of the year for each species. Yet, given the cryptic diversity of the tandem running behavior in non-neoisoptera termites, species-specific descriptive efforts are essential.

## Author contributions

N.M.: conceptualization, data curation, formal analysis, funding acquisition, investigation, methodology, project administration, validation, visualization, supervision, writing-original draft.

W.C.: formal analysis, methodology, writing-editing

E.C.: formal analysis, methodology, supervision, writing-editing T.N.: resources, writing-editing

T.K.: resources, writing-editing

## Data Availability Statement

All data and code used in this study are currently hosted on GitHub: https://github.com/williamchambliss/hodo-tandem. The repository will also be archived on Zenodo and assigned a DOI upon publication.

## Compliance with Ethical Standards

The authors declare no conflicts of interest. This research involved no human participants and did not require ethical approval for animal research.

## Acknowledgments

We thank Kensei Kikuchi for helping during the sampling of termites, Dr. Kenji Matsuura for the space for termite maintenance, and Dr. Thomas Bourguignon for providing experimental spaces. We acknowledge the use of ChatGPT, a language model developed by OpenAI, for minor suggestions with respect to the texts and coding. This study is supported by a JSPS (Japan Society for the Promotion of Science) Research Fellowship for Young Scientists CPD (Cross-border Post Doctorate) (20J00660) to N.M., a Grant-in-Aid for Early-Career Scientists (21K15168) to N.M., a Grant-in-Aid for Scientific Research B (22H02680) to T.K., IPSF fellowship from OIST to N.M., a Research grant for the Christopher Barnard Award from the ASAB (The Association for the Study of Animal Behaviour), and USDA National Institute of Food and Agriculture, Hatch projects number 7007938.

